# Spatiotemporal dynamics of multi-vesicular release is determined by heterogeneity of release sites in central synapses

**DOI:** 10.1101/2019.12.23.887372

**Authors:** Dario Maschi, Vitaly A. Klyachko

## Abstract

Synapses can release multiple vesicles in response to a single action potential. This multi-vesicular release (MVR) occurs at most synapses but its spatiotemporal properties and relation to uni-vesicular release (UVR) are poorly understood. Nanoscale-resolution detection of individual release events in hippocampal boutons revealed a pattern of spatial organization of MVR, which preferentially overlapped with UVR at more central release sites. Pairs of fusion events comprising MVR were also not perfectly synchronized and the earlier event within the pair occurred closer to the active zone (AZ) center. Parallel to this organization, individual release sites had a gradient of release probability extending from the AZ center to periphery. This gradient, and spatial features of MVR, were similarly tightened by buffering intracellular calcium. These observations revealed a heterogeneous landscape of release site properties within individual AZs, which determines the spatiotemporal features of MVR and is controlled by non-uniform calcium elevation across the AZ.

## INTRODUCTION

Information transmission in the brain relies critically on the number of vesicles released with each action potential, and major efforts have been made to model this process (Neher, 2010; Pan and Zucker, 2009; Rotman et al., 2011). Although initially it was hypothesized that, at most, only a single vesicle can be released from a given synapse with each action potential (i.e. univesicular release (UVR)), it is now widely accepted that two or more vesicles can fuse simultaneously in response to a single action potential in the same synaptic bouton, leading to the notion of multi-vesicular release (MVR) (Rudolph et al., 2015). MVR is a ubiquitous release mechanism in large and small central synapses throughout the brain (Auger et al., 1998; Christie and Jahr, 2006; Foster et al., 2005; Huang et al., 2010; Leitz and Kavalali, 2011; Malagon et al., 2016; Oertner et al., 2002; Rudolph et al., 2011; Singer et al., 2004; Taschenberger et al., 2002; Tong and Jahr, 1994; Wadiche and Jahr, 2001). It has been suggested to serve a wide range of functions including enhancing synaptic reliability, controlling synaptic integration, enhancing the efficient information transmission by complex spikes, and induction of several forms of plasticity (Rudolph et al., 2015). Despite its prevalence, spatiotemporal organization of MVR within the synaptic active zone (AZ) and its regulation are poorly understood. Investigating these questions has been hampered by the extremely small size of the AZ (Schikorski and Stevens, 1997, 1999), and limited resolution of conventional experimental approaches. As a result, it remains unknown whether all release sites within the AZ support both UVR and MVR, or specialized subsets of release sites are preferentially used for one form or the other. More generally, a fundamental unresolved question is whether all release sites are uniform in their properties across the AZ, or a more complex spatiotemporal landscape of release features exists within individual AZs and controls the occurrence of UVR and MVR.

We recently were able to overcome these limitations by developing a nanoscale imaging modality that enabled us to resolve the locations of individual vesicle fusion events with ∼27nm precision in active hippocampal synapses in culture (Maschi and Klyachko, 2017). These tools uncovered the presence of multiple distinct release sites within individual AZs, which undergo repeated rounds of re-use during UVR. Here we applied nanoscale imaging tools to detect and study organizational principles of MVR in individual hippocampal synapses in dissociated neuronal cultures. Our results revealed a non-random spatial and temporal organization of MVR events relative to the AZ center which parallels the spatial distribution of release site properties. The gradient of release site properties and the spatial features of MVR were also similarly affected by buffering intracellular calcium. Together our analyses suggest a new level of organization of the AZ. The spatiotemporal features of MVR reflect a heterogeneity of release site properties across individual AZs and depend, in part, on the non-uniform landscape of calcium elevation across the AZ following an action potential.

## RESULTS

### Heterogeneous Release Probability of Sites Supporting Uni- and Multi-Vesicular Release

To detect MVR events and resolve their locations in hippocampal synapses we took advantage of a nanoscale imaging approach we recently developed (Maschi and Klyachko, 2017) combined with the use of a pH-sensitive indicator pHluorin targeted to the vesicle lumen via vGlut1 (vGlut1-pHluorin) (Balaji and Ryan, 2007; Leitz and Kavalali, 2011; Voglmaier et al., 2006). vGlut1-pHluorin was expressed in cultures of excitatory hippocampal neurons using a lentiviral infection at DIV3 and imaging was performed at DIV 16-19 at 37°C. Release events were evoked using 1 Hz stimulation for 120 sec (**Figure 1A**). Robust detection of individual release events was achieved at 40 ms/frame rate throughout the observation time period. Using a hierarchical clustering algorithm, simulations and statistical considerations we have previously determined that release events are not randomly distributed throughout the AZ, but occur in a set of defined and repeatedly reused release sites within the AZs (Maschi and Klyachko, 2017) (see Methods for details).

**Figure 1:**
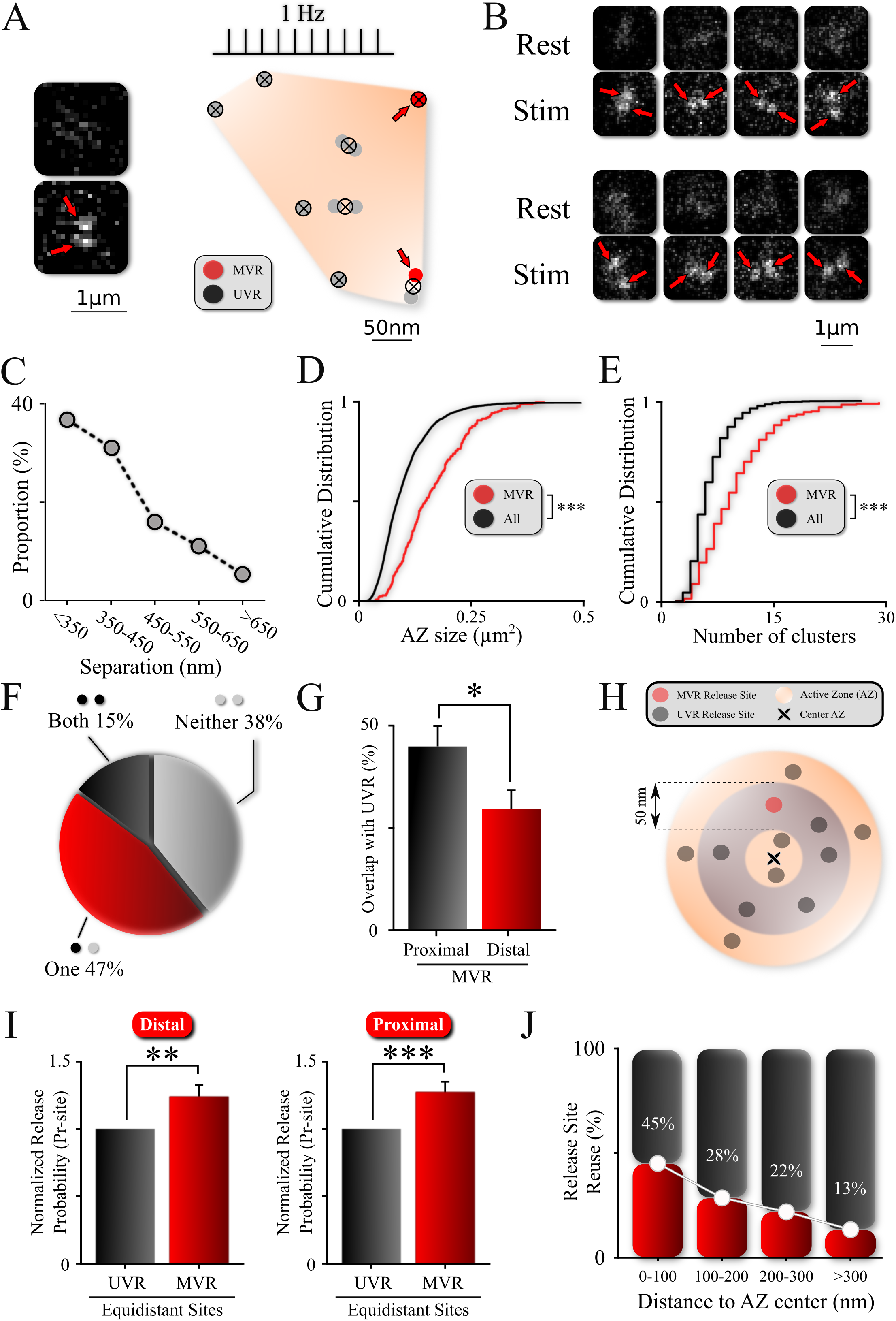
Non-Uniform Spatial Features of MVR Events and Release Sites within an AZ. **A**. Sample spatial distribution of ten UVR (grey) and MVR (red) events within a hippocampal bouton evoked by 120 APs at 1Hz. Cluster/release site locations are shown by crossed circles. Images (before and after 1 AP stimulation) show a sample MVR event highlighted by arrows. **B**. Examples of MVR events in different buttons. **C**. Proportion of MVR events as a function of intra-event separation distances. **D.E**. Cumulative distributions of AZ area (**D**) and number of clusters/release sites (**E**) for all recorded boutons (black) and boutons exhibiting MVR events (red). **F**. Spatial overlap of MVR and UVR events. Percentage of MVR events in which none, one or both events in the pair occurred at release sites that also harbored at least one UVR event. **G**. Probability of reuse by UVR events of more proximal vs. more distal release sites engaged in MVR event pairs. **H.I**. Analysis of release probability (P_r_-site) of more distal (**I**, left bars) and more proximal (**I**, right bars) release sites engaged in MVR event pairs compared to UVR events equidistant to the AZ center within ± 25nm (shown schematically in grey in **H**). **J**. Proportion of release sites that are reused at least once during observation period as a function of the distance to AZ center. Numbers shown represent average release site re-use in a given bin. N=3781(UVR); 245(MVR) events, from 90 dishes from 11 independent cultures; *p<0.05, **p<0.01, ***p<0.001. Two-sample KS-test(**D, E**); Chi-square test (**G**); Paired t-test (**I**).

Visual examination of these recordings revealed a subset of events with simultaneous fusion of two vesicles in the same bouton following a single action potential (**Figure 1A, B**). To automatically identify these dual events and determine the precise spatial locations of the two events comprising MVR, we used a well-established mixture-model fitting approach with two fixed-width Gaussians to approximate the PSF-like images of each vesicle (Jaqaman, 2008; Thomann et al., 2003). We previously used a conceptually similar fitting approach to localize individual release events with a ∼27nm precision for non-overlapping UVR events (Maschi and Klyachko, 2017). Here we found that MVR events occurred at a wide range of distances separating the event pair, but were more likely to occur at shorter separation distances (**Figure 1C**). Over ∼90% of MVR events had a separation distance below 600nm between the two events in the pair. We have previously used 3D FIB-SEM reconstruction of our neuronal cultures to show that the average bouton-to-bouton distance is an order of magnitude larger than event-to event distance and that the chances of misidentifying two events in the neighboring boutons as occurring in the same bouton is negligible (Maschi and Klyachko, 2017). We note that this mixture-model fitting approach does not reliably fit a subset of strongly overlapping MVR events that occur at very close distances; we thus examined this subset of unresolved MVR events using additional computational tools presented in the subsequent section.

We first asked how incidence of MVR is distributed. Previous studies suggested that release probability of a synapse is a strong predictor of a propensity for MVR (Christie and Jahr, 2006; Huang et al., 2010). It is also known that the AZ size is a major determinant of release probability in hippocampal synapses (Holderith et al., 2012; Matz et al., 2010). We thus explored the relationship between the AZ size (see Methods for details) and probability of observing MVR events at individual boutons. We also used the number of release sites per bouton as a functional measure of synapse release probability. Individual release sites within each bouton were defined using a hierarchical clustering algorithm with a cluster diameter of 50nm (Maschi and Klyachko, 2017). Boutons that exhibited MVR events had a significantly larger AZ (**Figure 1D** and **Table 1**; N=3781(UVR); 245(MVR) events, 90 dishes from 11 independent cultures; P<0.001, two-sample KS-test) and a significantly larger number of release sites than the synapse population overall (**Figure 1E; Table 1**). These results suggest that boutons with larger AZs (and higher overall release probability as given by a larger number of release sites) are more likely to produce MVR events.

**Table 1:**
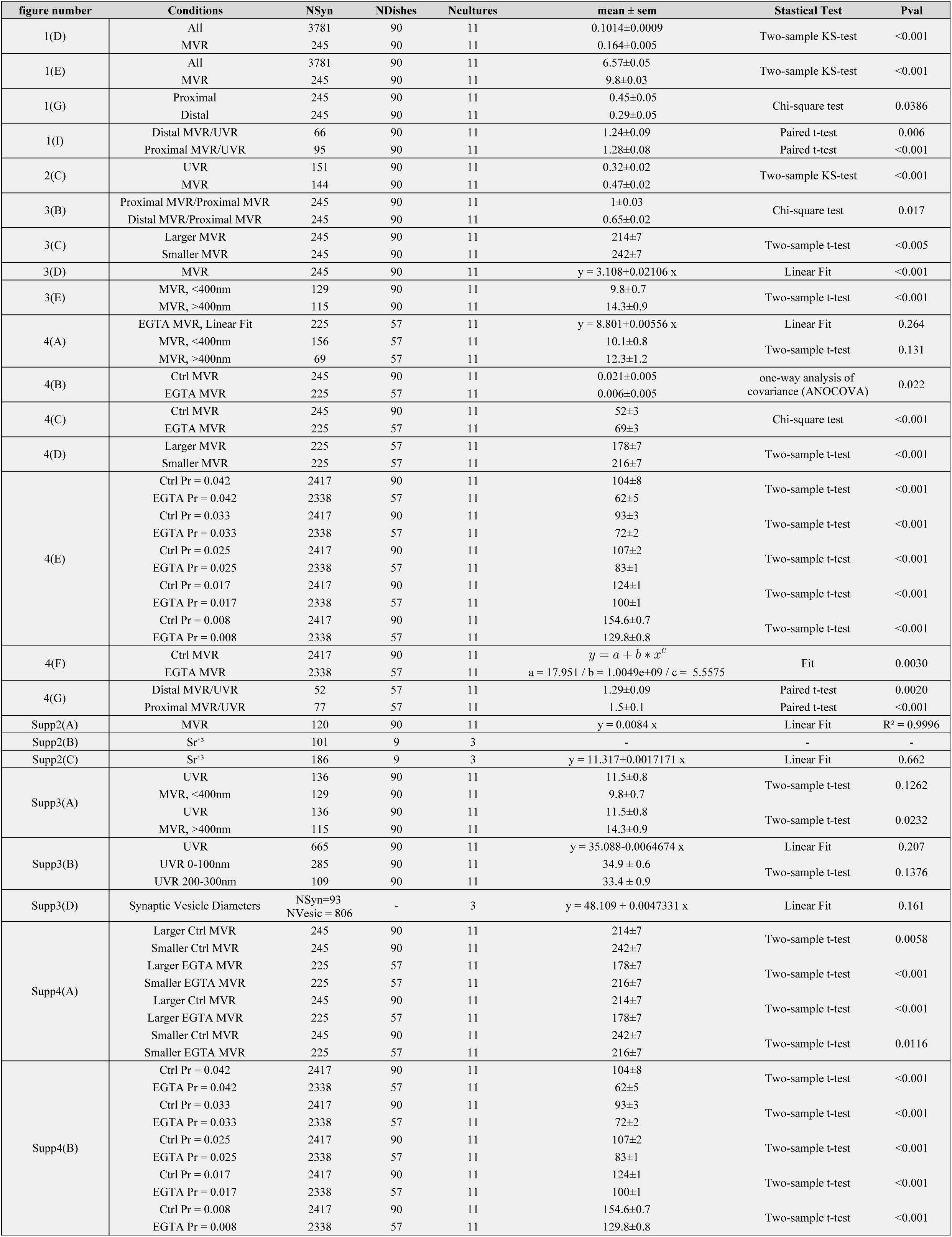
Data Values and Statistical Analyses. Columns represent (from left to right): figure/panel number; experimental conditions; number of samples (synapses, dishes, cultures); mean values and SEM; statistical test used for comparison; P-value resulting from the statistical comparison.

In addition to a variable propensity for MVR across the synapse population, we asked whether there is differential propensity for UVR versus MVR among release sites within the same bouton, or all release sites are equivalent in supporting both forms of release. We observed that MVR events were distributed throughout the AZ, but some heterogeneity could be seen between release sites engaged in UVR and MVR: there was no overlap with UVR for ∼38% of MVR events, a partial overlap in ∼47% (with one of the two events in the MVR pair overlapping with UVR); and a full overlap in ∼15% of MVR events (**Figure 1F; Table 1**). This result did not depend on the specific definition of release sites/clusters because we obtained essentially the same result using proximity analysis of individual events without using any clustering algorithms (**Figure S1; Table 1**). This observation suggests that release sites can harbor both UVR and MVR events, but are not uniform in their properties with respect to their capacity for UVR and MVR (explored in more detail below). We also note that because of a relatively short duration of observation, which is limited to 120 sec by a natural synapse displacement (Maschi and Klyachko, 2017), our results cannot be interpreted to indicate that there are some specialized release sites that *only* support MVR or UVR. Since MVR is a low probability event, much longer recordings would be necessary to determine if such sites exist.

To better understand why some sites may have higher propensity for MVR than others, we next examined whether UVR and MVR have distinct spatial patterns in the AZ plane. Specifically, for each of the two release sites engaged in the MVR pair, we determined the probability that the same release site is also re-used by UVR. We observed that the overlap of MVR and UVR events was location-dependent: the release site in the MVR pair that was more proximal to the AZ center had a significantly higher probability to be re-used by UVR events than the more distal one (**Figure 1G; Table 1**). Thus the spatial overlap between UVR and MVR events has a gradient from the AZ center to periphery, supporting the notion of heterogeneity of release site properties within the AZ.

To further examine this possibility, we quantified release probability at each release site (P_r_-site) based on the number of release events detected during the 120sec observation period. We then compared P_r_-site separately for each of the two events in the MVR pair with all other nearby release sites located equidistantly (i.e. within a ±25nm band) from the AZ center in the same bouton (**Figure 1H**). We observed that both release sites engaged in an MVR event had a significantly higher release probability than other equidistant UVR sites in the same bouton (**Figure 1I; Table 1**). More generally, we observed that release sites were highly heterogeneous in terms of release probability, which varied ∼5 fold among release sites within the same AZ (P_r_-site range [0.008-0.042]) during the observation time. Most importantly, we observed a spatial gradient of release probability among release sites, which decayed with distance from the AZ center (**Figure 1J**). These findings demonstrate a large degree of heterogeneity of release site properties within the individual AZs and suggest that release sites that have a higher propensity for MVR are characterized by a higher release probability than other sites equidistant to the AZ center.

### Spatiotemporal Features of Resolved MVR Events Generalize to Unresolved MVR Events

The well-separated MVR events analyzed thus far had a sufficient spatial separation for each event in the pair to be individually localized (resolved events). Because of the very small size of the AZ, a significant proportion of MVR events had sub-diffraction distances between the two events in the pair that could not be individually resolved. Therefore we ask to what extent the conclusions we obtained from the resolved MVR events generalize to the population of unresolved MVR events.

To identify unresolved MVR events we took advantage of the quantal nature of vesicular release to distinguish MVR from UVR events based on amplitude (Balaji and Ryan, 2007; Leitz and Kavalali, 2011). At 2mM extracellular Ca^+2^, over 90% of the events are UVR in hippocampal neurons at 37°C (Leitz and Kavalali, 2011; Maschi and Klyachko, 2017). We thus analyzed individually each synaptic bouton with a minimum of 5 fusion events to determine the mean and intrasynaptic variability (standard deviation) of quantal event amplitude. A threshold for the MVR event detection was set at two standard deviations above the mean quantal event amplitude determined individually for each bouton (**Figure 2A, B**). Based on this analysis, we estimated that such MVR events represent ∼9% of all release events; within this population, we could robustly separate ∼50% of the MVR from UVR events based solely on their amplitude (**Figure 2B**). We note that this approach to identify MVR events does not rely on spatial information, and thus is complementary to the mixture-model fitting approach we used above. Thus the populations of MVR events identified with these two approaches are partially overlapping (by ∼20%, data not shown).

**Figure 2:**
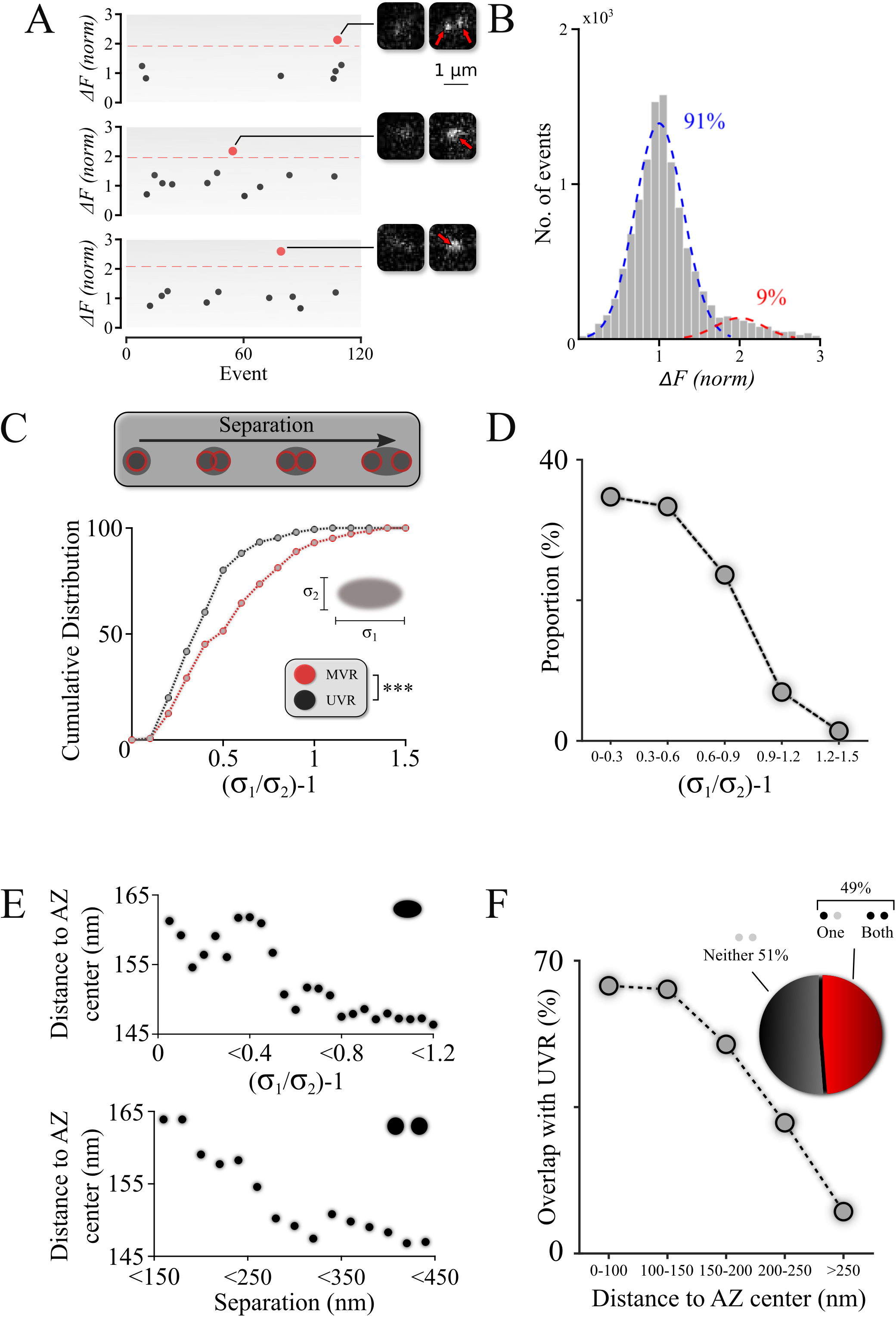
Spatiotemporal Features of Resolved MVR Events Generalize to Unresolved MVR Events. **A**. MVR (red) and UVR (black) events were separated based on the event amplitude. Examples of identified resolved MVR event (top) and two unresolved MVR events (middle and bottom) are shown with corresponding images. **B**. Intensity histogram for all detected events from (**A**) reflects the quantal nature of fusion events. Gaussian fits to the first peak (UVR, blue) and second peak (MVR, red) and their relative abundance are shown. **C**. Asymmetry analysis of unresolved MVR events vs UVR events. Asymmetry score was calculated using asymmetrical Gaussian fit to the event image to determine maximal (δ_1_) and minimal (δ_2_) width (*insert*). **D**. Proportion of unresolved MVR events as a function of asymmetry score which correlates with intra-event separation distances. **E**. Mean distance to the AZ center for unresolved (*top*) and resolved (*bottom*) MVR events. Distance was calculated from the peak of the Gaussian fit for unresolved MVR events and from the center of the line connecting two fusions within the resolved MVR events. **F**. Probability of overlap of unresolved MVR and UVR events in the same bouton as a function of distance to the AZ center. Event overlap was determined using proximity analysis with a radius of 25nm. Only more symmetrical MVR events (asymmetry score<0.5) were included in this analysis. Points represent proportion of MVR events within a given distance band that overlapped with UVR. *Pie chart:* Proportion of unresolved MVR events that overlap or not with UVR during observation period. N=151(UVR); 144(MVR) events, 90 dishes from 11 independent cultures. ***p<0.001; Two-sample KS-test (**C**).

The identified MVR events were then analyzed based on asymmetry considerations, using an asymmetric Gaussian model fit that takes into consideration the pixelated nature of the image to determine the width (sigma) of the Gaussian fit in the maximal (longitudinal, δ_1_) direction and minimal (transverse, δ_2_) direction. The ratio δ_1_/δ_2_ −1 (asymmetry score) represents an estimate of asymmetry of the double event image, which correlates with the distance between the two sub-diffraction events forming the image (DeCenzo et al., 2010). Distributions of asymmetry scores for the single and double events indicate that they represent two distinct populations (**Figure 2C; Table 1**) and thus validate our approach to robustly distinguish unresolved MVR from UVR events.

We then examined the spatiotemporal features of the unresolved MVR events. First, we observed that unresolved MVR events preferentially have smaller asymmetry scores (**Figure 2D**) and thus tend to occur at shorter separation distances, similarly to the resolved MVR events (**Figure 1C**). Next, we examined localization of unresolved MVR events relative to the AZ center/periphery. We observed that more asymmetrical (more spatially separated) events occurred closer to the AZ center, while symmetrical events tend to be more peripheral (**Figure 2E, top**). An equivalent calculation for the resolved MVR events (see figure legend for details) showed a very similar relationship (**Figure 2E, bottom**), supporting the notion that the two subpopulations of MVR events have similar spatial properties. Finally, we examined the spatial overlap between unresolved MVR events and UVR. Only a subpopulation of strongly overlapping MVR events (asymmetry score <0.5) were used in this analysis, because these highly symmetrical events could be well-approximated by a single symmetrical Gaussian fit, making this analysis comparable to that of the resolved MVR events. We observed a similar extent of spatial overlap with the UVR of both unresolved and resolved MVR events (**Figure 2F and 1F**), and the overlap also preferentially occurred closer to the AZ center for both types of MVR events (**Figure 2F and 1G**).

Taken together, our results suggest that unresolved MVR events have similar spatiotemporal features to the resolved MVR events. Our observations thus likely generalize across the entire population of MVR events.

### Temporal Desynchronization of Release Events Comprising MVR

Heterogeneity of release site properties poses a question of whether events comprising MVR are heterogeneous not only in the spatial but also in the temporal domain. Indeed, evidence for a temporal jitter within the MVR event doublets on a millisecond timescale (∼1-5ms) has been reported in both excitatory and inhibitory cerebellar synapses (Auger et al., 1998; Auger and Marty, 2000; Crowley et al., 2007; Malagon et al., 2016; Rudolph et al., 2011). Initial inspection of resolved MVR events suggested that one of the two events is often noticeably larger than the other (**Figure 3A**). Given that action potentials are synchronized with the beginning of the frame acquisition, and considering quantal nature of fusion events, we hypothesized that this difference in amplitude could reflect an imperfect synchronization within MVR pairs leading to smaller number of photons being acquired for the second event in the pair (**Figure 3A, top**). To investigate this possibility we examined if there is a systematic relationship between the amplitude difference and spatial organization of the double events comprising MVR. We observed that the larger amplitude event within the MVR pair was more likely to occur closer to the AZ center than the smaller one in the pair (**Figure 3B; Table 1**). Accordingly, the average distance to the AZ center was significantly shorter for the larger event than the smaller one in the pair (**Figure 3C; Table 1**). Most importantly, this amplitude difference was correlated with the distance between the two events within the pair (**Figure 3D,E**). We note that a component of the amplitude differences can arise from the uncertainty in determination of fusion event amplitude, which we estimated to be ∼10% (**Figure S3A**). Thus the uncertainty in our measurements can account for the amplitude differences of the closely-spaced MVR events, but not the far apart MVR events (**Figure S3A**). Moreover, the positive correlation between the amplitude difference and spatial separation of the two events comprising MVR cannot be explained by random noise or measurement uncertainty. We thus interpreted our results to indicate that the amplitude difference within MVR pairs at least in part reflects imperfect synchronization. Interestingly, our results indicate that the earlier event in the MVR pair preferentially occurs closer to the AZ center, while the second, delayed event occurs more peripherally. This spatial organization parallels the gradient of release site release probability from AZ center to periphery (**Figure 1J**, and see below).

**Figure 3.**
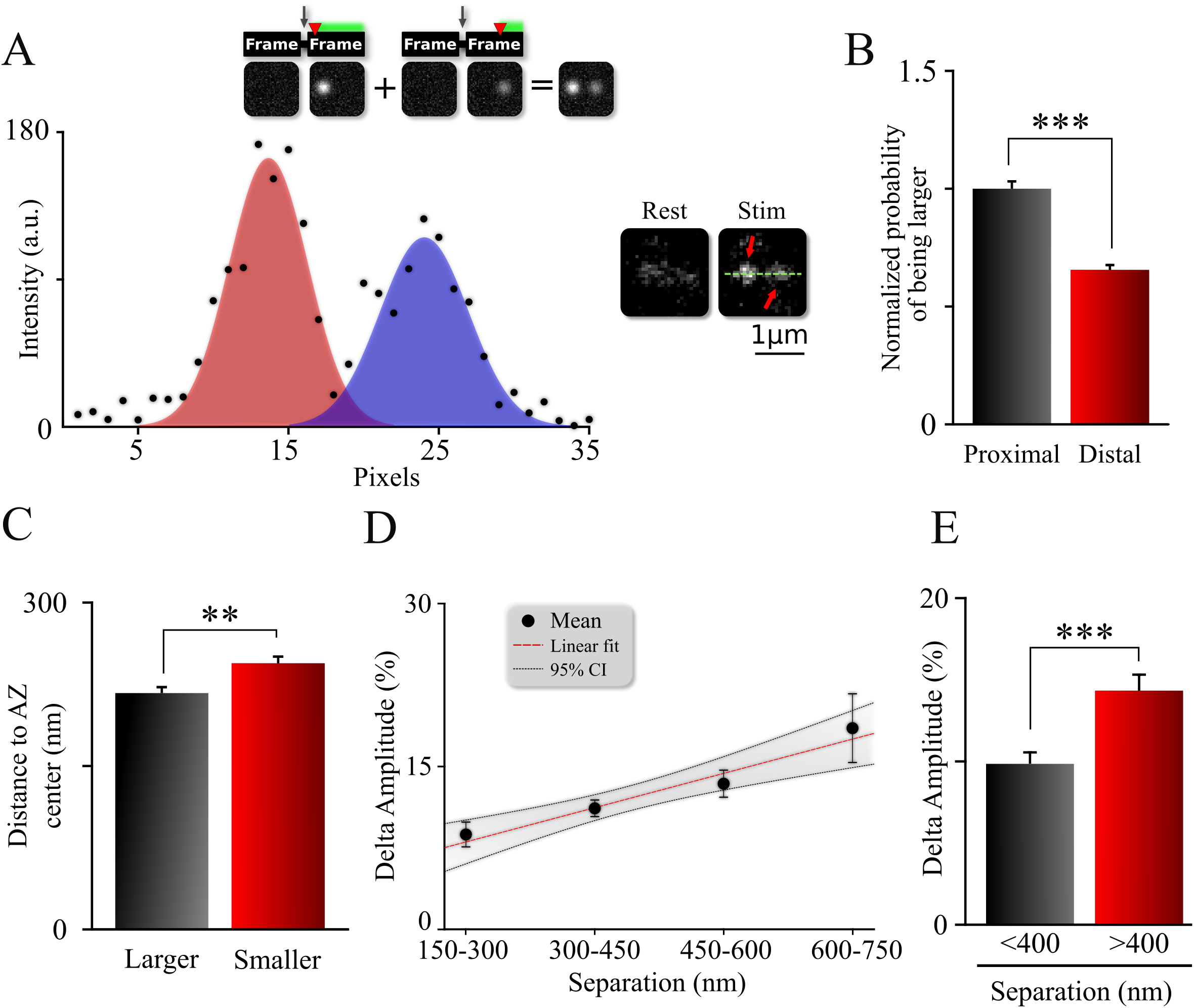
Spatiotemporal Organization of Release Events Comprising MVR. **A**. Sample image (*right*) and intensity profile (*left*) of an MVR event with noticeable difference in intra-event amplitude. Top *Insert* shows cartoon representation of a relationship between a time delay *(red arrow)* of the second fusion after an action potential and resulting amplitude difference within an MVR event. **B**. Probability to be larger for the proximal or distal event within the MVR pairs normalized to that of the proximal event. **C**. Distance to the AZ center from the larger and smaller events within MVR pairs. **D.E**. Amplitude difference of the two events comprising MVR as a function of intra-event separation. Linear fit (**D**) and t-test of pooled data (**E**) are shown. *p<0.05, ***p<0.001; Chi-square test (**B**); Paired t-test (**C**); Two-sample t-test (**E**). N=245(MVR) events, from 90 dishes from 11 independent cultures;

Given the observed amplitude differences within the MVR pairs, and the acquisition duration of 40ms per frame, we estimated the maximal time difference within double events in our recordings to be less than ∼4ms for the majority of events. We note that this value overestimates the true extent of desynchronization because a component of the differences arises from the uncertainty in the amplitude measurement. We estimated the maximal time delay within the MVR pairs to be ∼2ms, if the measurement uncertainty is factored in. These values are in a close agreement with the previous reports on desynchronization within MVR events, which was found to be in the range of 1-5ms (Auger et al., 1998; Auger and Marty, 2000; Crowley et al., 2007; Malagon et al., 2016; Rudolph et al., 2011).

Can factors other than desynchronization contribute to the difference in amplitude within MVR events and its spatial arrangement relative to the AZ center, such as differences in vesicle size or cleft pH along the AZ plane? We used the Large-Area Scanning Electron Microscopy (LaSEM) micrographs of our cultures (Maschi et al., 2018) to determine the size of all vesicles positioned next to the AZ (within 100nm, defined previously as tethered vesicles (Maschi et al., 2018a)) as a function of distance to the AZ center. Vesicle diameter did not exhibit any measurable changes with distance from the AZ center (**Figure S3C,D**). Thus the amplitude differences within MVR events are not due to systematic differences in vesicle size along the AZ. We further determined if the amplitude differences within MVR events can arise from differences in the cleft pH at different locales of the AZ. We examined the peak amplitude of vGlut1-pHluorin signal during individual fusion events, which is determined in a large part by the cleft pH, as a function of distance to AZ center. We found no measurable differences in the vGlut1-pHluorin signal amplitude (**Figure S3B**), suggesting that a gradient of cleft pH is unlikely to explain the differences in MVR event amplitude. Finally, we note that in our imaging experiments dozens of synapses are positioned in random orientation and are recorded simultaneously. Some boutons have the center of the AZ in focus and others have the periphery, and yet others are positioned in all other possible configurations in-between, all in the same recording. Thus there is no bias towards any one possible configuration or tilt of the AZ relative to the imaging plane that can lead to systematically larger fusion event amplitude at the AZ center vs. periphery. Indeed, this notion is highlighted by the fact that the average UVR event amplitude is indistinguishable at the AZ center vs periphery in our measurements (**Figure S3B, *insert***). Thus the observed differences in MVR event amplitude are unlikely to result from a bias in AZ position relative to the imaging plane.

In summary, our results support the notion of desynchronization within the MVR events and revealed a pattern of spatial organization of MVR in which the first of the two events in the MVR pair is preferentially located closer to the AZ center. This spatial organization of MVR parallels the spatial gradient of the release site properties within the AZs.

### Double events do not result from asynchronous release overlapping temporally with synchronous events

Given the intrinsically limited temporal resolution of our imaging tools, we asked whether the observed double events could arise not from MVR, but rather from asynchronous release events generated by preceding stimulation and temporarily overlapping with UVR. We considered this possibility in five complementary ways:

First, if our double events arise from overlap of synchronous release with asynchronous events generated by preceding stimulation, probability of such double events is expected to increase during the train. In contrast, we detected no increase in the double event probability during the stimulus train (**Figure S2A, Table 1**). In fact, probability of observing double events is slightly higher on the first stimulus for which there is no preceding stimulation. Thus, uniform probability of observing double events during trains argues against major contribution from asynchronous release.

Second, synchronous release events, including MVR, are time-locked with the stimulus and are acquired for the entire duration of the frame. Thus the amplitude distribution of MVR events should appear as a single Gaussian peak centered at ∼2q value (twice the amplitude of a UVR event), as indeed is the case in our measurements (**Figure 2B**). In contrast, asynchronous release by definition is not time-locked with the stimulus and thus occurs randomly at any time during the acquisition frame. As a result, single asynchronous release events are acquired for a wide range of durations which are less than a full frame, and thus must have a skewed non-Gaussian amplitude distribution shifted towards smaller values than a full size synchronous event. We confirm that this is the case using asynchronous events we recorded in 3mM Sr^2+^ in otherwise identical conditions (**Figure S2B, Table 1**). Thus our double events are synchronous events and have properties distinct from asynchronous release.

Third, we compared spatiotemporal properties of our double events with that of asynchronous release events we recorded in 3mM Sr^2+^. The asynchronous release events detected in the same frame in the same bouton (**Figure S2C**) did not exhibit the spatiotemporal features that we observed for the double events (**Figure 3D**). Thus spatiotemporal properties of the observed MVR events are distinct from properties exhibited by asynchronous release.

Forth, the high synchronicity of double events in our recordings argues against significant contribution from asynchronous release. As mentioned above, the estimated synchronicity within a few milliseconds in our double events is a good agreement with the previous measurements of MVR and agrees with an accepted definition of MVR (Auger et al., 1998; Auger and Marty, 2000; Crowley et al., 2007; Malagon et al., 2016; Rudolph et al., 2011).

Finally, we note that previous literature found minimal asynchronous release evoked by 1Hz stimulation under nearly identical experimental conditions (37°C, 2mM extracellular Ca^2+^) (Raingo et al, 2012).

In summary, five complimentary lines of evidence strongly suggest that double events in our recordings indeed represent synchronous MVR events, and not asynchronous release.

### Spatiotemporal Features of MVR events and Release Site Properties are Calcium-Dependent

What is the mechanistic origin of the spatial organization of MVR events relative to the AZ center? Our previous study suggested a possible role of calcium in spatial regulation of release site properties because we observed that release site usage during UVR is regulated in an activity-dependent manner with a shift of site usage from the AZ center toward the periphery during trains of activity (Maschi and Klyachko, 2017). Previous studies have also found that propensity for MVR scales with increase in extracellular calcium levels (Leitz and Kavalali, 2011). Whether calcium regulates the spatiotemporal organization of MVR and/or the properties of the release sites remains unknown. To test this possibility, we pre-incubated neurons with a cell-permeable calcium chelator EGTA-AM (30µm) for 20 min. EGTA does not directly affect vesicle release because it is too slow to buffer rapid calcium rise near VGCCs, but is effective in buffering the ensuing slower calcium elevation due to diffusion. EGTA had several effects on spatiotemporal features of MVR events. First, EGTA affected the MVR event desynchronization: while the amplitudes within the double events were still different, the difference no longer depended on the distance between the two events (**Figure 4A,B; Table 1;** N= 225 synapses, from 57 dishes from 11 independent cultures). Second, the spatial organization of the MVR events was also affected by EGTA with a larger proportion of MVR events occurring at shorter intra-event distances in the presence of EGTA (**Figure 4C; Table 1**). Accordingly, the average distances from both events in the MVR pair to the AZ center were significantly shortened in the presence of EGTA (**Figure S4A, Table 1**). Thus calcium buffering causes MVR events to occur at shorter separation distances and more proximal to the AZ center. However, the preferential localization of the earlier event in the MVR pair closer to the AZ center than the second, delayed event was still observed in the presence of EGTA (**Figure 4D, Table 1**). These results suggest that several major spatiotemporal features of MVR are determined, in part, by calcium diffusion following an action potential.

**Figure 4:**
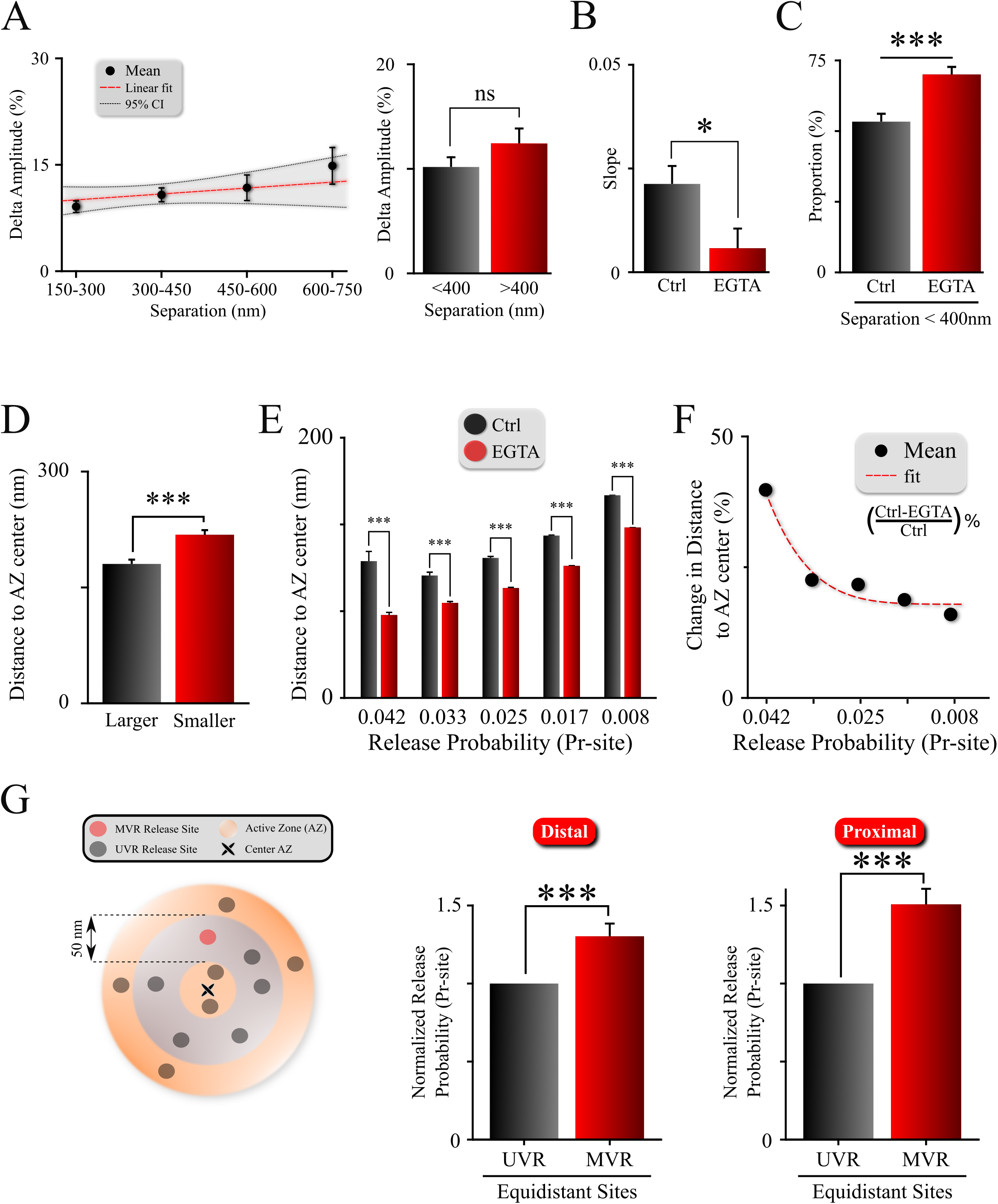
Spatiotemporal Features of MVR Events and Release Site Properties are Calcium-Dependent. **A**. Effect of EGTA on a correlation between the spatial separation and amplitude difference between two events comprising MVR. **B**. Effect of EGTA was assessed by comparing the slopes of the correlation in (A), in control (from Figure 3D) and EGTA (from Figure 4A) conditions. **C**. Proportion of MVR events with intra-event separation <400 nm in EGTA and control conditions. **D**. Distance to the AZ center from the larger and smaller events within MVR pairs in the presence of EGTA. **E.F**. Average distance to the AZ center (**E**) and its relative change (**F**) of individual release sites binned based on their release probability, in EGTA and control conditions. **G**. Release probability of more distal (left bars) and more central (right bars) release sites engaged in MVR event pairs compared to all other release sites equidistant to the AZ center within ± 25nm, in the presence of EGTA. ***p<0.001; ns, not significant. Two-sample t-test (**A,E);** one-way analysis of covariance (ANOCOVA); Chi-square test (**C**); Paired t-test (**D,G**). Control: N=245 MVR events from 90 dishes from 11 independent cultures. EGTA: N=225 MVR events from 57 dishes from 11 independent cultures.

In line with the idea that spatiotemporal features of MVR events reflect release site heterogeneity, we observed that preferential utilization of more central release sites was exacerbated in the presence of EGTA **(Figure 4E, S4B; Table 1)**. Interestingly, EGTA affected stronger, more central release sites to a larger extent than the weaker, more peripheral ones (**Figure 4F**). These results are consistent with our earlier findings that an opposite change in the release site utilization/reuse from AZ center towards periphery occurs during high-frequency stimulation (Maschi and Klyachko, 2017). It is also consistent with the shorter separation distance within MVR events in the presence of EGTA (**Figure 4C**), and shorter distances of MVR events to the AZ center in the presence of EGTA (**Figure S4A**). Indeed, MVR events tend to occur at release sites with higher release probability, the utilization of which shifts closer to the AZ center in the presence of EGTA, and the preferential usage of higher release probability sites by MVR events (as compared to equidistant UVR sites) persisted in the presence of EGTA (**Figure 4G; Table 1**).

Together these observations suggest that the gradient of release site properties as well as spatiotemporal features of MVR events are, in part, determined by the landscape of calcium rise across the AZ.

## DISCUSSION

While MVR is well established as a ubiquitous release mechanism in many types of synapses (Rudolph et al., 2015), its spatiotemporal organization and regulation at the AZ is largely unknown. We took advantage of nanoscale detection of individual release events in central synapses to reveal three major findings: (i) MVR events exhibit spatial and temporal patterns of organization. MVR preferentially occurs at release sites with higher release probability and is more likely to overlap with UVR closer to the AZ center. MVR events are also not perfectly synchronized and spatially organized with the first of the two events located closer to the AZ center. (ii) Parallel to this organization of MVR, release sites within the same AZ have highly heterogeneous properties with a gradient of release probability from the AZ center to periphery. (iii) The gradient of release site properties as well as spatiotemporal features of MVR, are determined, in part, by the intraterminal calcium elevation following an action potential. Together these results suggest that non-uniform spatiotemporal dynamics of MVR arises from heterogeneity of release site properties within the individual AZs. Our results suggest a model of MVR in which the earlier of the two events in the MVR pair is similar to a UVR event in that it occurs closer to the AZ center because release sites with higher release probability are localized preferentially more proximally to the AZ center. A second release event is then triggered occasionally with a short delay after the same action potential, at a more peripheral release site primed for release, in part due to calcium spread from the AZ center. This notion is supported by effects of calcium buffering with EGTA and with our observations that both events in the MVR pair occur at sites with higher release probability than other equidistant sites in the same bouton. These observations reveal a previously unrecognized complex landscape of release probability within the AZ, which determines the spatial propensity for MVR and arise, in part, from a gradient of calcium across individual AZs.

What are molecular underpinnings of release site heterogeneity? Recent nanoscopy studies indicate that release sites colocalize with nanoclusters of presynaptic docking factors, such as RIM1/2 (Tang et al., 2016), which have been suggested to control recruitment and clustering of VGCCs at the AZ via the RIM binding protein RIM-BP (Davydova et al., 2014; Hibino et al., 2002; Wang et al., 2000). RIM1/2 nanoclusters are more likely to be located near the center of the AZ than in the periphery, suggesting a possible structural basis for the gradient of release site properties (Tang et al., 2016). Moreover, enrichment of many scaffold/docking proteins within their clusters is dynamic, including RIM, Bassoon, and Munc13 (Bademosi et al., 2016; Glebov et al., 2018; Smyth et al., 2013; Tang et al., 2016; Weyhersmüller et al., 2011). Thus the heterogeneity of release site properties may arise, in part, from variability in cluster architecture, such as cluster size or relative enrichment of tethering/docking/priming factors. An additional source of heterogeneity could arise from the fact that variable fractions of many critical components of release machinery, including VGCCs, Syntaxin-1, and Munc18, are mobile within the AZ (Glebov et al., 2018; Schneider et al., 2015; Smyth et al., 2013). For example, a large proportion (>50%) of VGCCs are mobile in the AZ plane and this mobility is calcium-dependent (Schneider et al., 2015). How VGCC mobility is spatially controlled has not been explored, but differential VGCC stability at the AZ center vs periphery could account, in part, for the heterogeneity of the calcium rise across the bouton. VGCC mobility could also affect coupling between the channels and the vesicles (Eggermann et al., 2011; Miki et al., 2017), which could explain the differential effect of EGTA we observed on peripheral vs central release sites. Another possibility is that release site properties are determined, in part, by extrinsic factors. For example, release site refilling and vesicle retention at release sites depends on actin and myosins (Maschi et al., 2018; Miki et al., 2016). Thus a non-homogeneous spatial distribution of actin cytoskeleton could contribute to differential release site usage across a bouton. In addition, calcium influx at least through some subtypes of VGCCs is also modulated by the balance of phosphorylation/dephosphorylation by CDK5 and calcineurin (Kim and Ryan, 2013), and is directly regulated by a number of presynaptic proteins, such as ELKS (Liu et al., 2014), and Munc13 (Calloway et al., 2015). While the factors driving release site heterogeneity remain to be elucidated, our results uncovered a previously unknown level of structural and functional organization of release sites within individual AZs that determines the spatiotemporal dynamics of MVR.

## EXPERIMENTAL METHODS

### Neuronal cell cultures

Neuronal cultures were produced from rat hippocampus as previously described (Peng et al., 2012). Briefly, hippocampi were dissected from E16 pups, dissociated by papain digestion, and plated on coated glass coverslips. Neurons were cultured in Neurobasal media supplemented with B27. All animal procedures conformed to the guidelines approved by the Washington University Animal Studies Committee.

### Experimental Design

All measurements were replicated in more than 100 boutons derived from 22-90 coverslips from 7-11 independent cultures (see **Table 1** for individual experiments). Most experiments were carried out in an unblended manner and a specific randomization strategy was not used. Statistical computations were not performed to determine the optimal sample size for experiments.

### Lentiviral Infection

VGlut1-pHluorin was generously provided by Drs. Robert Edwards and Susan Voglmaier (UCSF) (Voglmaier et al., 2006). Lentiviral vectors were generated by the Viral Vectors Core at Washington University. Hippocampal neuronal cultures were infected at DIV3 as previously described (Maschi and Klyachko 2017).

### Fluorescence Microscopy

All experiments were conducted at 37°C within a whole-microscope incubator (In Vivo Scientific) at DIV16–19. Neurons were perfused with bath solution (125 mM NaCl, 2.5 mM KCl, 2 mM CaCl_2_, 1 mM MgCl_2_, 10 mM HEPES, 15 mM Glucose, 50 μM DL-AP5, 10 μM CNQX adjusted to pH 7.4). Asynchronous release events were recorded using the same solutions except 3mM Sr^2+^ and 0 mM CaCl_2_ were used in the bath. Fluorescence was excited with a Lambda XL lamp (Sutter Instrument) through a 100x 1.45 NA oil-immersion objective and captured with a cooled CMOS camera (Hamamatsu). With this configuration the effective pixel size was 65 nm. Focal plane was continuously monitored, and focal drift was automatically adjusted with ∼10 nm accuracy by an automated feedback focus control system (Ludl Electronics). Field stimulation was performed by using a pair of platinum electrodes and controlled by the software via Master-9 stimulus generator (AMPI). Images were acquired using an acquisition time of 40ms, one 45ms before stimulation and one coincidently (0ms delay) with stimulation.

### Large-Area Scanning Electron Microscopy (LaSEM)

LASEM methods and data used were published previously (Maschi et al, 2018). Briefly, cells were grown on 12 mm glass coverslips, were aspirated of media and fixed in a solution containing 2.5% glutaraldehyde and 2% paraformaldehyde in 0.15 M cacodylate buffer with 2 mM CaCl2, pH 7.4 that had been warmed to 37°C for one hour. The samples were then stained according the methods described by Deerinck et. al. 2010. Large areas (∼ 330 × 330 µm) were then imaged at high resolution in a FE-SEM (Zeiss Merlin, Oberkochen, Germany) using the ATLAS (Fibics, Ottowa, Canada) scan engine to tile large regions of interest. High-resolution tiles were captured at 16,384 × 16,384 pixels at 5 nm/pixel with a 5 µs dwell time and line average of 2. The SEM was operated at 8 KeV and 900 pA using the solid-state backscatter detector. Tiles were aligned and export using ATLAS 5.

### Image and Data Analysis

#### Event detection and localization

The fusion event localization at subpixel resolution was performed using custom-written Matlab code based on the uTrack software package that was kindly made available by Dr. Gaudenz Danuser lab (Aguet et al., 2013; Jaqaman, 2008). The input parameters for the PSF were determined using stationary green fluorescent 40 nm beads.

Localization precision was determined directly from least-squares Gaussian fits of individual events as described in (Thomann et al., 2003; Thomann et al., 2002) using in-built functions in uTrack software (Aguet et al., 2013; Jaqaman, 2008). Spatial constraints of the vesicle lumen imply that only a few VGluT-pHluorin molecules can be located within individual vesicle. Given our observations that the fluorescence signal evoked by vesicle fusion did not disperse significantly during our acquisition time of 40ms, the small number of VGluT-pHluorin molecules per vesicle and their lateral movement upon fusion, if present, do not strongly affect localization precision at the time our measurements are made.

Localization of resolved MVR events (Figures 1,3 and 4) was performed using a mixture-model multi-Gaussian fit as described in (Thomann et al., 2003; Thomann et al., 2002) using in-built functions in uTrack software (Aguet et al., 2013; Jaqaman, 2008).

Unresolved MVR events (Figure 2) were identified based on the event amplitude. The single event amplitude and its variability were determined for each bouton individually. Photobleaching was accounted for by fitting the event intensity changes over time. The threshold for MVR event detection was set at two standard deviations above the mean single event amplitude determined individually for each bouton. Localization of unresolved MVR events was determined using an asymmetrical Gaussian model fit based on the minimization of the residuals.

#### Definition of release sites

Release sites were defined using hierarchical clustering algorithm using built-in functions in Matlab as described (Maschi and Klyachko 2017; Maschi et al. 2018; Wang et al., 2016). We have previously compared the results of this clustering analysis obtained with experimentally observed distribution of fusion events vs. the same number of simulated events distributed randomly across the same AZs (Maschi and Klyachko, 2017). We found that randomly distributed release events result in a very different pattern of clustering than experimentally observed events, and do not reproduce the observed features of real release event clusters. The observed clusters thus do not arise from random distribution of release events, but rather represent a set of defined and repeatedly reused fusion sites within the AZs.

#### Release site release probability

Release probability of individual release sites was calculated based on the number of release events detected per release site and divided by the duration of the observation period.

#### AZ dimensions and center

The AZ size was approximated based on the convex hull encompassing all vesicle fusion events in a given bouton. This measurement is in a close agreement with the ultrastructural measurements of AZ dimensions (Maschi and Klyachko, 2017). AZ center was defined as the mean position of all fusion events in a given bouton.

#### Event proximity analysis

To determine overlap of MVR and UVR events, a proximity analysis was performed in which overlap was defined as having at least one UVR event occurring within 25nm of an MVR event during observation period.

#### Synapse identification and analysis of vesicle diameter in LaSEM data

Three characteristic features were used for synapse identification: the presence of a synaptic vesicle cluster, the postsynaptic density and the uniform gap between pre and postsynaptic membranes. Distance to the AZ center was determined from the projection of vesicle center position to the AZ plane.

#### Fit regression models

Nonlinear and linear fit regression models were generated using built-in functions in Matlab.

### Data Inclusion and Exclusion Criteria

A minimum of 5 detected release events per bouton was required for all analyses.

### Statistical Analysis

Statistical analyses were performed in Matlab. Statistical significance was determined using two-sample two-tailed t-test, paired t-test, Kolmogorov-Smirnov (K-S) test, one-way analysis of covariance (ANOCOVA) or chi-square test, where appropriate. The number of experiments reported reflects the number of different cell cultures tested. The value of N is provided in the corresponding figure legends and **Table 1**. Statistical tests used to measure significance are indicated in each figure legend along with the corresponding significance level (p value). Data are reported as mean ± SEM and p < 0.05 was considered statistically significant. Analysis of the samples was not blinded to condition. Randomization and sample size determination strategies are not applicable to this study and were not performed.

## Acknowledgements

This work was supported in part by the R35 grant to VAK from NINDS. We thank Dr. Meyer Jackson for helpful comments on the manuscript.

## SUPPLEMENTARY FIGURE LEGENDS

**Supplementary Figure 1:**
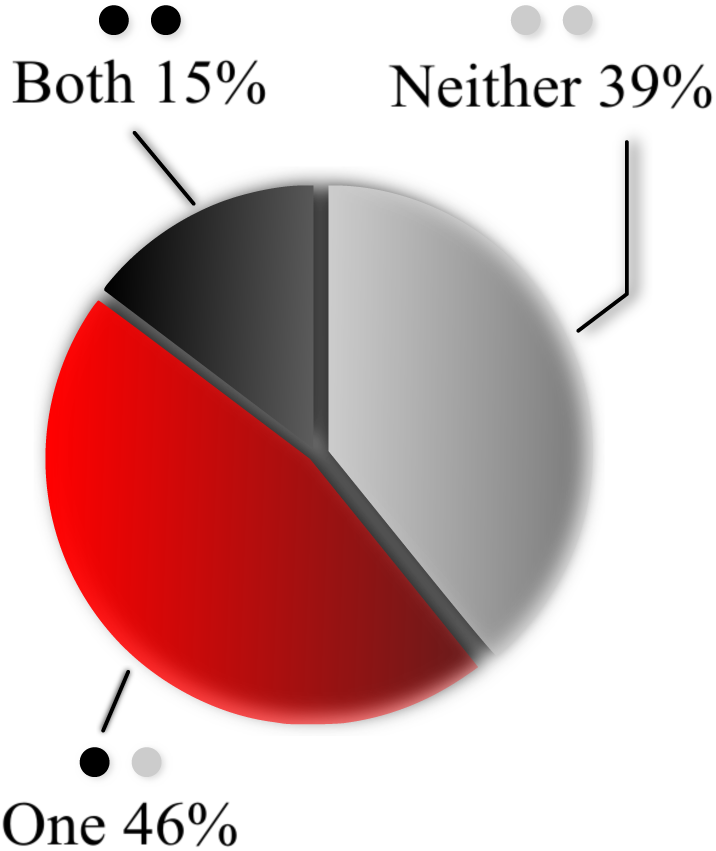
Overlap of MVR and UVR events determined by proximity analysis. Percentage of MVR events in which none, one or both events in the pair overlapped (within 25nm) with at least one UVR event during observation period, as determined by proximity analysis.

**Supplementary Figure 2:**
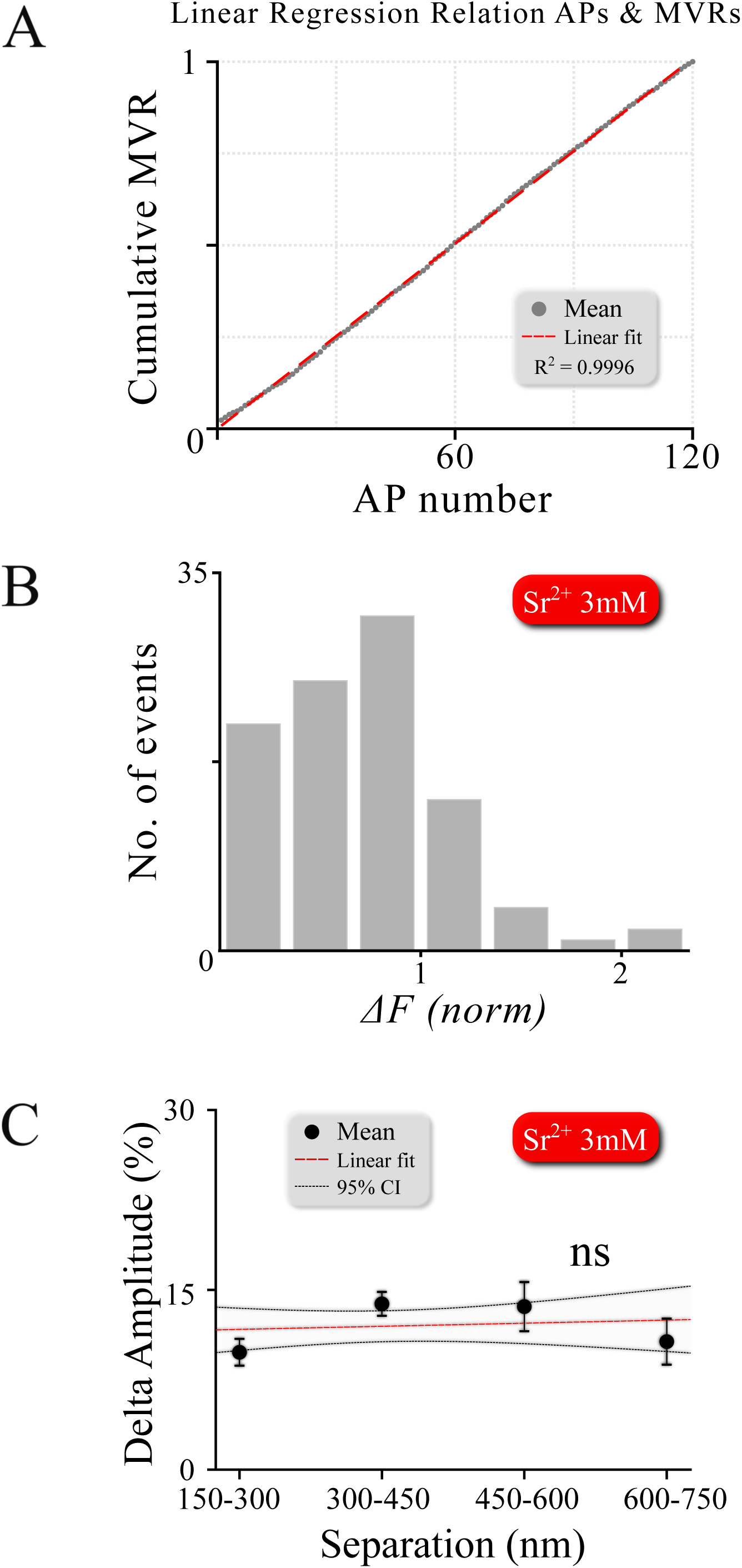
Double events do not result from asynchronous release overlapping temporally with synchronous events. **A**. MVR events occur at the same rate throughout the 120 action potential stimulus at 1Hz, showing absolutely no increase in double event probability during the train, as would be expected if these events are generated by asynchronous release. **B**. Amplitude histogram for asynchronous release events detected in 3mM Sr^+2^ after single APs using 1Hz 120 AP stimulation shows the loss of quantal homogeneity of asynchronous fusion events (with a left-shifted non-Gaussian distribution). This is distinct from the Gaussian amplitude distribution we observed for synchronous uni- and multi-vesicular events in our measurements. **C**. Amplitude difference of two asynchronous events detected in the same frame in the same bouton in the presence of 3mM Sr^2+^ as a function of intra-event separation. In contrast to MVR, no correlation was found between spatial event separation and desynchronization (p = 0.662).

**Supplementary Figure 3:**
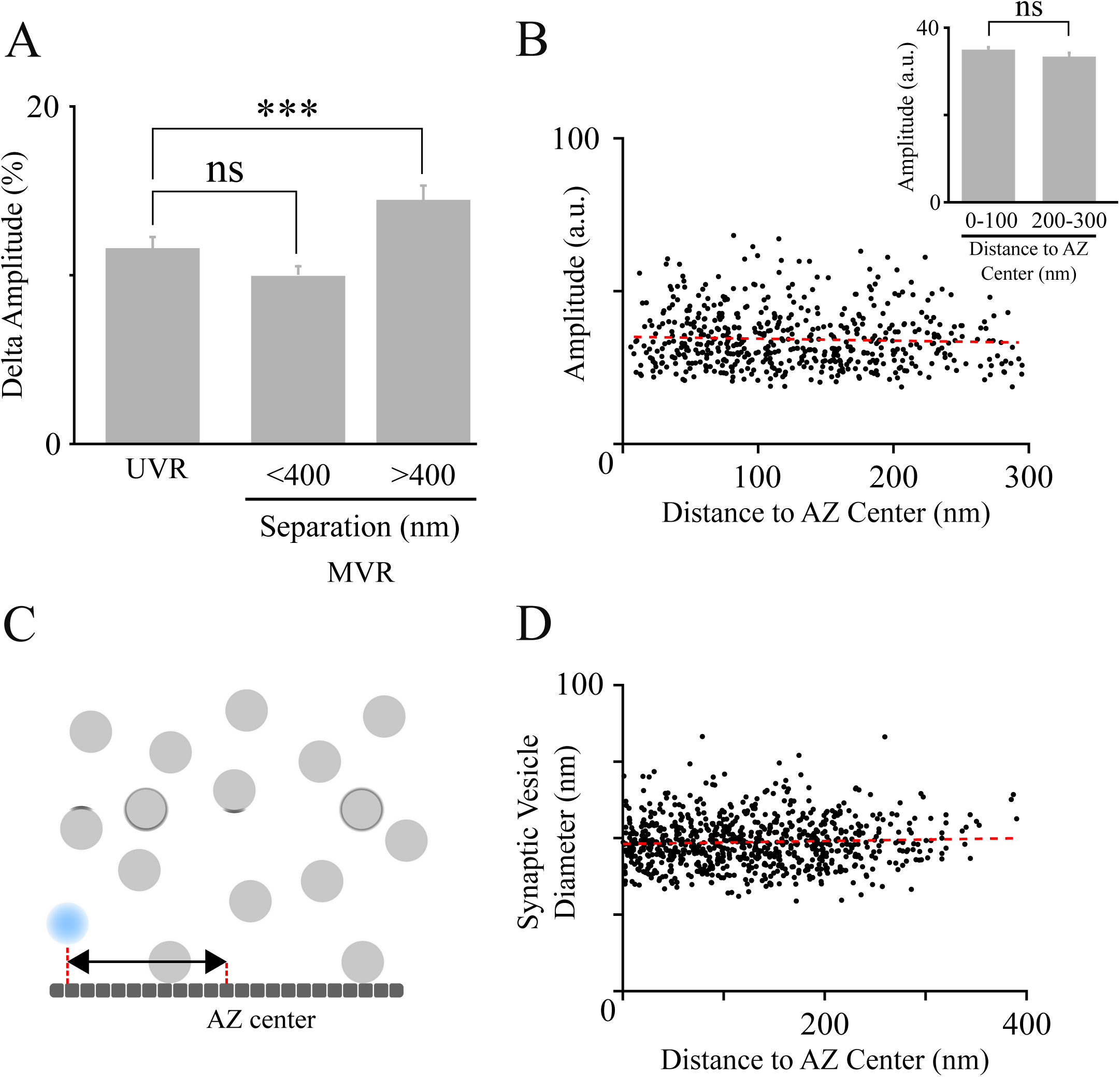
Amplitude difference within the MVR event pairs is not due to measurement uncertainty, changes in vesicle size or cleft pH within the AZs. **A**. Uncertainty in event amplitude determination was estimated based on the amplitude variation of consecutive UVR events evoked in the same bouton at 1Hz. Uncertainty in determination of individual event amplitude does not account for the amplitude differences observed in MVR events with large separation. **B**. Amplitude of UVR events as a function of the distance to the AZ center. No correlation was found between event amplitude and distance to AZ center (p = 0.21, linear fit). Bar graph shows no significant difference in amplitude exists between central events (pooled data for 0-100nm) vs. peripheral events (pooled data for 200-300nm). **C.D**. Vesicle diameter as a function of distance to the AZ center was determined for all vesicles positioned within 100nm from the AZ (defined previously as tethered vesicles (Maschi et al., 2018)) from LaSEM micrographs of hippocampal cultures (EM data from Maschi et al., 2018). Distance from vesicle to AZ center was determined from the projection of the vesicle center position onto the AZ plane (C). Vesicle diameter did not show significant changes with distance from the AZ center (p = 0.16, linear fit). ***p<0.001; ns, not significant. Two-sample t-test (**A,B)**.

**Supplementary Figure 4:**
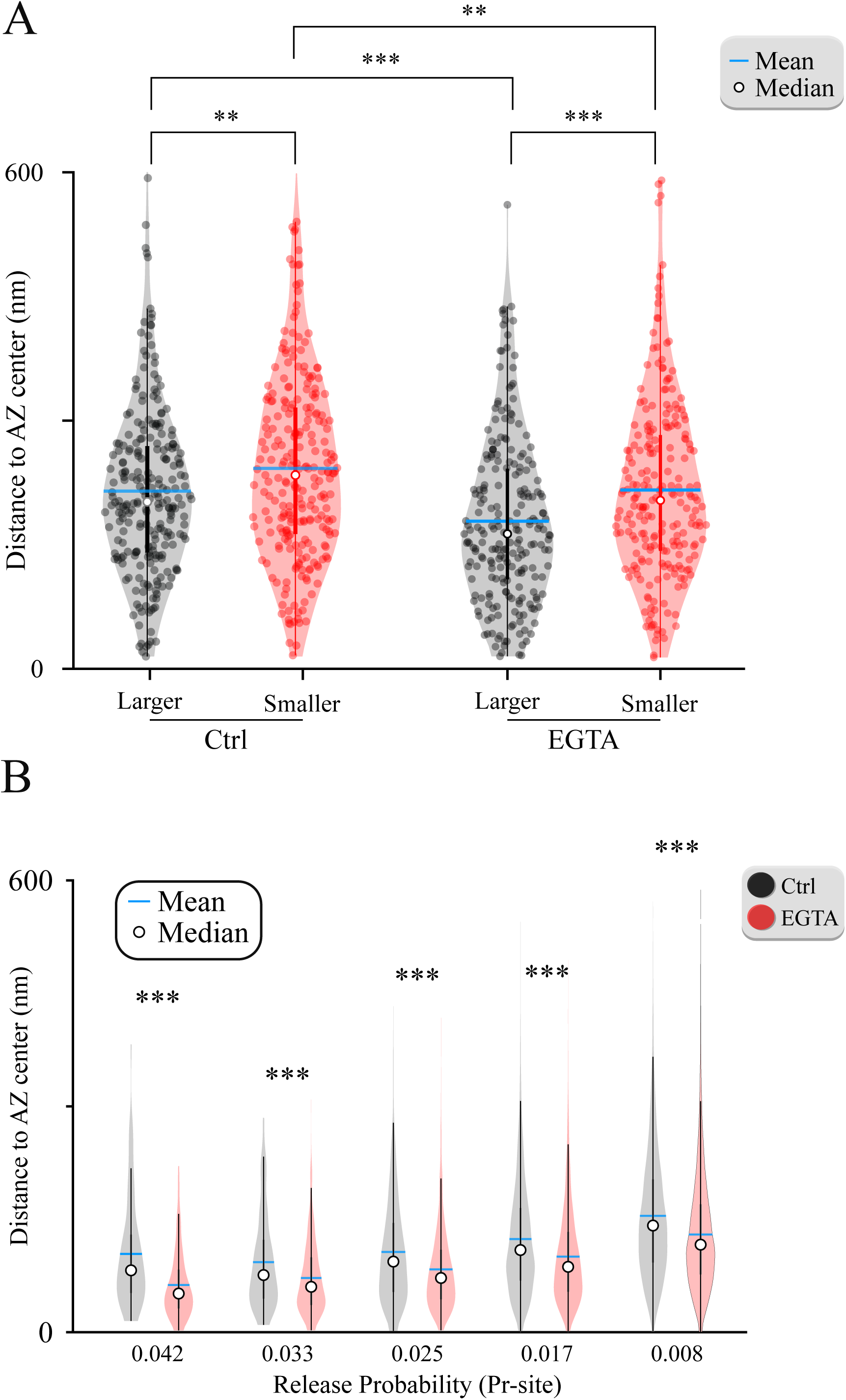
Violin plots of the amplitude differences within MVR event pairs and the effects of EGTA. **A**. Distance to the AZ center from the larger and smaller events within MVR pairs in control (data from **Figure 3C**) and EGTA conditions (data from **Figure 4D**). **B**. Average distance to the AZ center of individual release sites binned based on their release probability, in control and EGTA conditions (data from **Figure 4E**). **p<0.01; ***p<0.001. Two-sample t-test.

